# Thymus Involution across the Human Menstrual Cycle

**DOI:** 10.64898/2026.07.15.738838

**Authors:** Julian Koenig, Holger Winkels, Tobias Sodenkamp, Kimberly E. Schroeder, Jessica C. Sieren

## Abstract

**Importance:** Thymic involution in humans has traditionally been conceptualized as a linear, age-associated process. Animal studies suggest transient hormone-dependent fluctuations in thymus size during reproductive transitions, including across the estrous cycle, but translational evidence in humans remains limited.

**Objective:** To determine whether thymus size changes across the menstrual cycle in healthy women and whether these changes are associated with fluctuations in estradiol and progesterone levels.

**Design:** Observational repeated-measures study using quantitative computed tomography (qCT). Data were collected as part of an imaging study examining menstrual cycle–related physiological variation. Participants underwent assessments during menses and the early luteal phase within the same menstrual cycle. Mixed-effects models with robust variance estimation evaluated associations between cycle phase, hormonal contraception, reproductive hormone levels, and thymus size. Thymus segmentation was independently conducted by 2 blinded raters.

**Setting:** Single-center university-based imaging study conducted at the University of Iowa.

**Participants:** Thirty-one non-smoking women with regular menstrual cycles were included, of whom n = 16 used oral hormonal birth control and n = 15 did not use hormonal contraception. Exclusion criteria included pregnancy, breastfeeding, postmenopausal status, diabetes, body mass index greater than 30 kg/m^2^, hysterectomy, or use of long-term noncyclic hormonal contraception. Participants tracked menstrual cycles using temperature monitoring and ovulation kits before completing visits during menses and the early luteal phase. All participants provided informed consent before study participation.

**Main Outcomes and Measures:** Primary outcome was thymus size measured in mm^3^ using ultra-low-dose qCT imaging. Secondary outcomes included serum estradiol and progesterone levels.

**Results:** Estradiol levels increased from menses to the early luteal phase independent of hormonal contraceptive status. Progesterone levels were significantly lower among women using hormonal birth control. Thymus size differed significantly by hormonal contraceptive group, with larger thymus volumes observed among women not using hormonal birth control. Among women not using hormonal contraception, 66.7% demonstrated thymic involution from menses to the early luteal phase, compared with 37.5% of women using hormonal contraception. Estradiol and progesterone significantly interacted in predicting thymus size.

**Conclusions and Relevance:** These findings provide first-in-human evidence suggesting that thymus involution demonstrates short-term dynamics across the menstrual cycle and may be influenced by reproductive hormones and hormonal contraception.

**Key Points:** *Question:* Does thymus size change dynamically across the human menstrual cycle in association with fluctuations in reproductive hormones and hormonal contraception use?

*Findings:* In this repeated-measures quantitative computed tomography study of n = 31 healthy women, thymus size demonstrated within-person variation across the menstrual cycle. Greater thymic involution was associated with larger increases in estradiol, whereas increases in thymus size were associated with larger increases in progesterone; thymus size also differed significantly by hormonal contraceptive status.

*Meaning:* These findings suggest that thymic involution in humans may demonstrate short-term hormone-dependent dynamics beyond age-related atrophy, with potential implications for sex differences in immune regulation and inflammatory disease.

## Introduction

Thymus involution in humans is considered a linear process associated with aging. Thymic size has been shown to shrink at an annual rate of approximately 3% per year until 40 years of age and at an annual rate of 1% thereafter.^1^ Mice show transient changes in thymus size during phases of hormonal transition. Early studies illustrated pregnancy-induced involution of the thymus in mice,^2^ even when controlling for changes in body weight.^3^ Following studies identified in pregnant mice inversely correlated thymus weight with progesterone levels. Importantly, the association between thymus weight and progesterone is observed progressively over pregnancy in mice,^4^ and the pregnancy-induced decrease in thymus size is transient, recovering 2-weeks postpartum.^5^ No such association is present in non-pregnant and pseudo-pregnant mice.^6^ Animal studies across the estrous cycle further illustrate that the proestrus stage - associated with a peak in circulating estrogen – is characterized by the greatest decrease in thymus weight.^7^ Early castration in cattle, pigs and rabbits, has been shown to delay thymus involution.^8^ Chemical and surgical castration even reversed age-related thymic involution in rats.^9^ Experimental studies further illustrate dose dependent effects of sex-steroid administration on thymus size in rats and mice, with higher doses of estrogen administration resulting in a greater decrease of thymus weight.^10^ Data on dynamic within-subject changes in thymus size during phases of hormonal transition in humans are scarce. One study^11^ reported on two pregnant patients with signs of pulmonary embolism who underwent repeated computer tomography (CT) imaging. Both cases showed an enlarged thymus decreasing in size following birth. Changes in thymus size across the menstrual cycle in humans have previously not been studied. We hypothesized that thymus size demonstrates dynamic within-subject variation across the menstrual cycle in humans, with greater thymic involution associated with increases in estradiol and modulation by progesterone and hormonal contraceptive use.

## Methods

We drew on existing data from quantitative CT (qCT) in n = 31 non-smoking, healthy women with regular menstrual cycles (n = 16 on cyclic hormonal birth control), who underwent qCT imaging at menses (M) and the early luteal phase (ELP).^12^ Subjects were recruited from campus advertisements and normal, non-smoking cohorts from other research studies examining the lungs. Exclusion criteria included subjects who were pregnant, breastfeeding, body mass index > 30 kg/cm2, post-menopausal, diabetic, were taking long-term non-cyclic birth control (i.e., implant, patch, or injected), or had a hysterectomy. All study subjects provided written informed consent to participate in the study and data was collected with approval from the University of Iowa Institutional Review Board (IRB# 201606790), in accordance with the Declaration of Helsinki.

Women were asked to track their menstrual cycle via temperature and an ovulation kit for 1-2 months, before completing two study visits (M and ELP). The two study visits were completed within the same month/cycle for most subjects and were completed over more than one cycle in some cases due to scheduling restrictions. At each visit, subjects had peripheral blood collected and underwent ultra-low-dose qCT imaging. Hormone levels of estradiol and progesterone were analyzed from peripheral blood using the Roche Diagnostics Estradiol III and Progesterone III assays processed by the Emory Warner Pathology Laboratories.^13^ Ultra-low-dose, non-contrast chest qCT imaging was performed using a SOMATOM Force CT scanner (Siemens Healthcare) with 120 kV, 36 ref mAs, CareDose on, pitch 1.2, 0.5-s rotation time, 0.5-mm slice thickness, and Qr40 (admire 3, < 5-s scan). The CTDI_vol_ for one scan at 36 mAs was approximately 2.4 mGy or 1.0 mSv. Thymus size in mm3 was manually traced in ITK-SNAP,^14^ independently by two raters (TS and KES). Raters were blinded to group and phase and showed good agreement (two-way mixed ICC 0.895; 95%: [0.823;0.936]). The average of both raters was used for all analyses.

Data were analyzed using mixed models with robust variance estimation, addressing the main effect of cycle phase (M/ELP) and group (hormonal birth control/no hormonal birth control) as well as their interaction on hormone levels and thymus size, using the subject ID as random factor over repeated measures. Contrasts were Bonferroni corrected and all analyses were adjusted for age and days between repeated assessments. The relative change in thymus size was described descriptively in change in percent from M to ELP. Additional mixed models predicted the change in thymus size based on hormone interactions. Simple linear regression was used to illustrate bivariate associations. Statistical analyses were performed with Stata/SE (version 18.0; StataCorp LP, College Station, TX, US), with a set significance level of α = 0.05.

## Results

Females not on hormonal birth control (NBC) were older compared to those on oral hormonal birth control (OBC; t_(29)_ = 2.779, *p <*.*01*). Full descriptive data by group and cycle phase is provided in *Table 1*. Estradiol showed a significant main effect of cycle phase (model: χ^2^ _(62/31)_ = 19.84, *p =*.*001;* phase: χ^2^ = 6.15, *p =*.*013*), suggesting a significant increase from M to the ELP independent of group (NBC/OBC). No significant effect of group nor a significant group by phase interaction was observed. Progesterone showed a significant main effect of group (model: χ^2^ _(62/31)_ = 16.11, *p =*.*01;* group: χ^2^ = 9.83, *p <*.*01*), suggesting lower progesterone in females OBC in contrast to those NBC. No significant effect of cycle phase nor a significant group by phase interaction was observed. Individual trajectories in estradiol and progesterone by phase of the menstrual cycle are illustrated in *Figure 1A*. Thymus size (mm^3^) showed a significant main effect of group (model: χ^2^ _(62/31)_ = 19.68, *p =*.*001;* group: χ^2^ = 6.93, *p =*.*01*), suggesting greater thymus size in NBC in contrast to OBC. No significant effect of phase nor a significant group by phase interaction was observed. Individual trajectories of thymus size by phase of the menstrual cycle are illustrated in *Figure 1B*. In NBC, n = 10 (66.67%) showed thymus involution from M to ELP (−0.81 to -30.32% change in relative size) while thymus size increased in n = 5 (+2.98 to +66.59% change in relative size). In OBC, n = 6 (37.50%) showed thymus involution from M to ELP (−5.81 to -28.43% change in relative size) and n = 10 showed an increase in thymus size (+9.69 to +55.34% change in relative size). Estradiol and progesterone showed a significant interaction in predicting thymus size: χ^2^ _(62/31)_ = 53.76, *p <*.*001;* interaction: coef. = -6.85, *p < 0*.*01*. As illustrated in *Figure 1C*, greater thymus involution, independent of group, was associated with a greater increase in estradiol from M to ELP, whereas an increase in thymus size was associated with a greater increase in progesterone from M to ELP.

**Table 1:** Descriptive data by group and phase of the menstrual cycle; M: menses; ELP: early luteal phase; SD: standard deviation; DRSP: drospirenone; DSG: desogestrel; NGM: norgestimate; LNG: levonorgestrel; EE: ethinyl estradiol; *: n = 6 assays in the OBC group below the measurement threshold of <5 pg/ml, imputed with 5 pg/ml; **: n = 2 assays in the OBC group below the measurement threshold of <0.2 ng/mL, imputed with 0.20 ng/ml

|  | no birth control (NBC) |  | on birth control (OBC) |  |
| --- | --- | --- | --- | --- |
|  | *M* | *ELP* | *M* | *ELP* |
| n | 15 |  | 16 |  |
| Age in years, mean (SD),<br>[range] | 31.93 (9.48)<br>[21-51] |  | 24.63 (4.44)<br>[20-38] |  |
| Days between visits, mean<br>(SD), [range] | 24.07 (21.12)<br>[7-77] |  | 19.94 (11.34)<br>[9-44] |  |
| Type/dosing of birth<br>control, n (%) |  |  |  |  |
| NGM-EE | 0 (0.00) |  | 6 (37.50) |  |
| DRSP-EE | 0 (0.00) |  | 8 (50.0) |  |
| LNG-EE | 0 (0.00) |  | 1 (6.25) |  |
| DSG-EE | 0 (0.00) |  | 1 (6.25) |  |
| Dosing in mg, mean (SD),<br>[range]* | 0 (0.00)<br>[0.00-0.00] |  | 0.03 (0.01)<br>[0.02-0.035] |  |
| Estradiol in pg/mL, mean<br>(SD), [range]* | 49.60<br>(43.09)<br>[6-178] | 97.73<br>(62.74)<br>[21-207] | 32.44<br>(26.54)<br>[5-91] | 102.50<br>(183.05)<br>[5-560] |
| Progesterone in ng/mL,<br>mean (SD), [range]** | 1.65 (2.79)<br>[0.20-10] | 3.51 (4.09)<br>[0.20-10.70] | 0.36 (0.17)<br>[0.20-0.90] | 0.41 (0.16)<br>[0.20-0.70] |
| Thymus size (mm<sup>3</sup>), mean<br>(SD), [range] | 19269.28<br>(9035.55)<br>[9757.17-38414.95] | 18573.06<br>(8745.54)<br>[7629.94-36998.55] | 13428.74<br>(4050.06)<br>[4424.41-20231.40] | 14940.71<br>(6392.18)<br>[6872.69-28805.90] |

**Figure 1:**
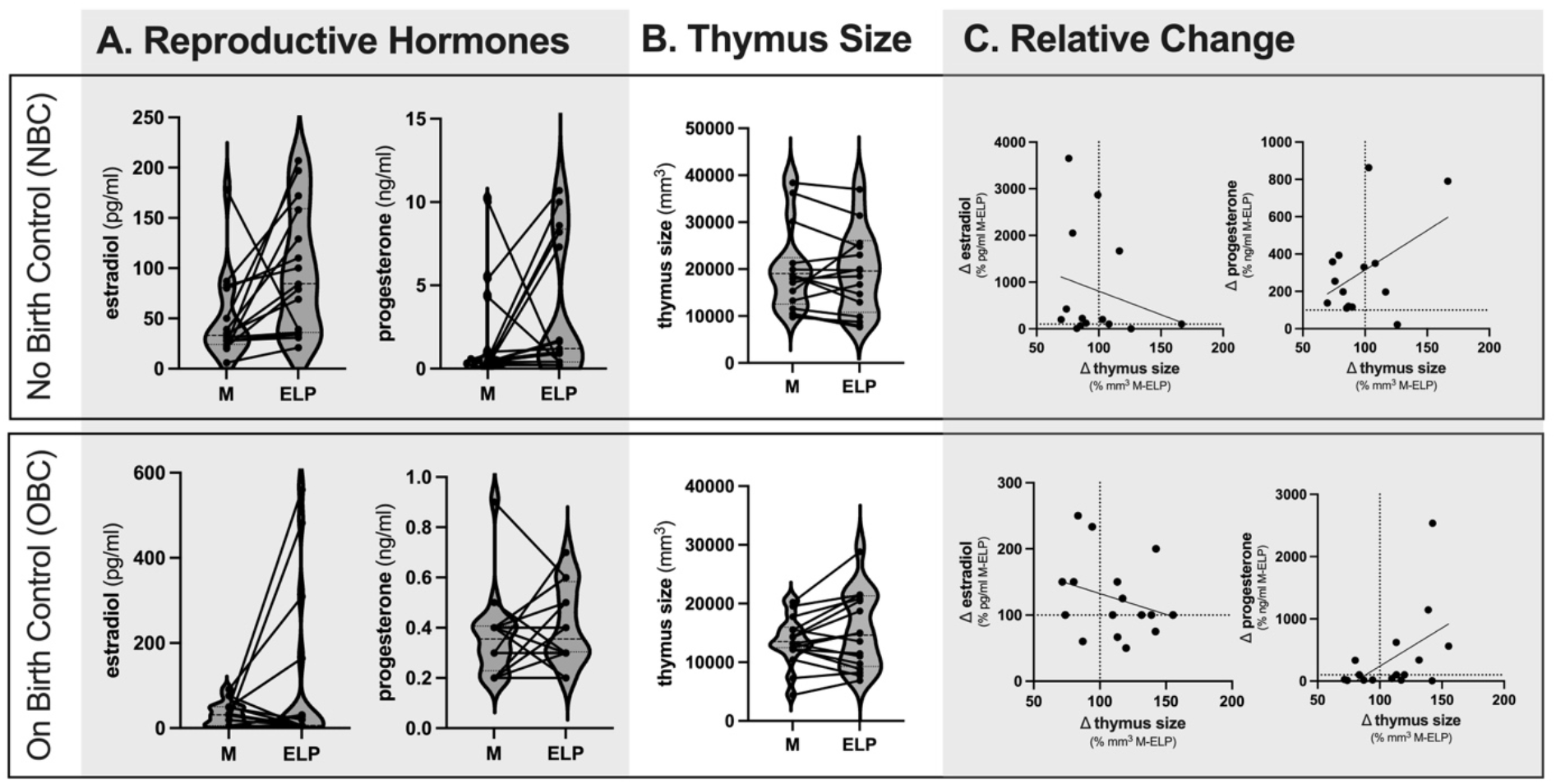
Individual Change in Reproductive Hormones and Thymus Size by Group and Phase; M: menses; ELP: early luteal phase;

## Discussion

To our knowledge, the present data are first-in-human, showing secondary within-subject dynamics of thymus size on the level of days (not years). Thymus size shows intra-individual variance across the menstrual cycle and is further influenced by hormonal contraception. Our findings challenge established conceptions of thymus involution in humans. The functional significance of these findings, however, remain speculative. Thymic atrophy in aging has been suggested to contribute to immunosenescence and inflammaging via decreased thymic output of naïve T cells.^15^ Here we illustrate secondary dynamics of thymus involution as a function of fluctuating reproductive hormones and hormonal contraception, with widespread implications for female health across the lifespan.

Only recently, thymectomy has been shown to reduce new T-cell production and health in adulthood,^16^ suggesting that the thymus remains functionally important beyond puberty. A general decrease in T lymphocytes is observed during the menstrual period in humans.^17^ Specifically, CD4+ T cells are significantly lower in the midluteal phase compared to the early follicular phase.^18^ Studies addressing hormone-dependent changes in thymus size in association with thymocyte release and composition of recent thymic emigrants are warranted. Dynamic changes in thymus size across the menstrual cycle may explain corresponding fluctuations in pro-inflammatory markers,^19,20^ via a secondary arc to the cholinergic inflammatory reflex, sensitive to changes in reproductive hormones. Efferent signaling of the vagus nerve innervates with macrophages in the periphery, releasing acetylcholine (ACh) which inhibits the synthesis of pro-inflammatory cytokines.^21^ The cholinergic anti-inflammatory pathway, modulates via α7 nicotinic acetylcholine receptor (α7nAChR) macrophage inflammation by inhibiting HMGB1 and cytokine release.^22^ A non-neural source of ACh are T cells, of which only CD4+ T cells express choline acetyltransferase (ChAT), responsible for the synthesis of ACh. Splenic-sympathetic fibers are thought to release norepinephrine acting on ChAT CD4+ T cells, to release ACh, inhibiting tumor necrosis factor alpha (TNFα) released in splenic macrophages.^23^ ACh thereby attenuates the release of IL-6 and other cytokines.

Decreased availability of ChAT expressing CD4+ T cells following dynamic thymic involution may thus compromise the cholinergic anti-inflammatory reflex and control of low-grade inflammation during phases of hormonal transition, explaining greater risk for inflammatory disorders in females.

Challenging the dominance of a unisex view on physiological function, the suggested mechanism may explain sex disparities in the pathophysiology and response to treatment in a host of diseases. The further discovery of the proposed pathway may inform a new understanding of differential susceptibility to immune-mediated disorders (i.e., autoimmune thyroiditis, rheumatoid arthritis and multiple sclerosis), and could explain increased risk for mood and behavioral changes during phases of hormonal transition in females. Inflammation is a key pathogenetic factor in depression^24^ with elevated CRP and IL-6 observed among depressed females (not males).^25^ Both inflammatory markers (IL-6 and CRP) are thought to promote depression characteristic sickness behavior^26^ via central nervous system circuitry, and are linked to specific symptom profiles, including anhedonia and reduced motivation but also a loss of appetite and decreased concentration.^27^ Phases of hormonal transition (i.e., puberty, pregnancy, menopause) across the female lifespan are associated with an increased risk of depression onset, and females also show variance in depressed mood and suicidal ideation across the menstrual cycle,^28^ leading to the formulation of distinct psychiatric entities, including the Premenstrual Syndrome (PMS) and Premenstrual Dysphoric Disorder (PMDD). Dynamic involution of the thymus in the presence of changing reproductive hormones, may be present as contributing pathogenic factor, triggering an inflammatory cascade in female depression.

The present study has several limitations that need to be addressed. qCT imaging is limited in the reliable assessment of thymus size. Our approach to manual tracing showed reasonable inter-rater agreement, dual energy CT imaging with iodinated contrast enhancement or magnetic resonance imaging and the use of neural network models for automated segmentation may lead to more robust results.^29^ Here we drew on secondary data from a small sample. Future prospective studies are warranted, including more frequent assessments over the menstrual cycle (and other phases of hormonal transition) to study trajectories in the association between sex hormones and thymus size at greater temporal resolution. Further, studies in men may provide important insights to general dynamics in thymus size independent of fluctuating reproductive hormones. Respective studies should further include the assessment of pro-inflammatory markers and lymphocyte composition, to address the functional significance of changes in thymus structure. Further, studies suggest compensatory mechanisms, illustrating an enlarged spleen after thymectomy and vice versa. Assessing spleen function and size alongside dynamics of the thymus during phases of hormonal transition seems warranted.

In conclusion, we present first in-human data, suggesting secondary dynamics to thymus involution, demonstrated here across the menstrual cycle. Findings challenge the prevailing textbook view of singular aging-associated thymus atrophy. The concept of thymus involution in humans is based on cross-sectional (predominantly post-mortem) samples, supporting a static view on thymus size (and function). Advances in neuroimaging technology enable to overcome these long-held beliefs and rethink the nature of the thymus.

## Declarations

### Human Ethics and Consent to Participate

All study subjects provided written informed consent to participate in the study and data was collected with approval from the University of Iowa Institutional Review Board (IRB# 201606790), in accordance with the Declaration of Helsinki.

### Data Sharing Statement

Research data are not shared due to legal and privacy issues.

## References

1. Pido-Lopez J, Imami N, Aspinall R. Both age and gender affect thymic output: more recent thymic migrants in females than males as they age. Clin Exp Immunol. 2001;125(3):409–413. doi:10.1046/j.1365-2249.2001.01640.x

2. Endo T, Kanayama K. Changes in the weight of the thymus after birth and in pregnancy in mice. Res Commun Mol Pathol Pharmacol. 1998;101(3):307–310.

3. Persike EC. Involution of Thymus During Pregnancy in Young Mice. Proceedings of the Society for Experimental Biology and Medicine. 1940;45(1):315–317. doi:10.3181/00379727-45-11666

4. Clarke AG, Kendall MD. The thymus in pregnancy: the interplay of neural, endocrine and immune influences. Immunol Today. 1994;15(11):545–551. doi:10.1016/0167-5699(94)90212-7

5. Rijhsinghani AG, Bhatia SK, Tygrett LT, Waldschmidt TJ. Effect of pregnancy on thymic T cell development. Am J Reprod Immunol. 1996;35(6):523–528. doi:10.1111/j.1600-0897.1996.tb00052.x

6. Chambers SP, Clarke AG. Measurement of thymus weight, lumbar node weight and progesterone levels in syngeneically pregnant, allogeneically pregnant, and pseudopregnant mice. J Reprod Fertil. 1979;55(2):309–315. doi:10.1530/jrf.0.0550309

7. Lee H, Kim H, Chung Y, Kim J, Yang H. Thymocyte Differentiation is Regulated by a Change in Estradiol Levels during the Estrous Cycle in Mouse. Dev Reprod. 2013;17(4):441–449. doi:10.12717/DR.2013.17.4.441

8. Henderson J. On the relationship of the thymus to the sexual organs: I. The influence of castration on the thymus. J Physiol. 1904;31(3-4):222–229. doi:10.1113/jphysiol.1904.sp001032

9. Kendall MD, Fitzpatrick FT, Greenstein BD, Khoylou F, Safieh B, Hamblin A. Reversal of ageing changes in the thymus of rats by chemical or surgical castration. Cell Tissue Res. 1990;261(3):555–564. doi:10.1007/BF00313535

10. Kuhl H, Gross M, Schneider M, et al. The effect of sex steroids and hormonal contraceptives upon thymus and spleen on intact female rats. Contraception. 1983;28(6):587–601. doi:10.1016/0010-7824(83)90109-9

11. Swami S, Tong I, Bilodeau CC, Bourjeily G. Thymic involution in pregnancy: a universal finding? Obstet Med. 2012;5(3):130–132. doi:10.1258/om.2011.110077

12. Sieren JC, Schroeder KE, Guo J, Asosingh K, Erzurum S, Hoffman EA. Menstrual cycle impacts lung structure measures derived from quantitative computed tomography. Eur Radiol. 2022;32(5):2883–2890. doi:10.1007/s00330-021-08404-9

13. Lippe BM, LaFranchi SH, Lavin N, Parlow A, Coyotupa J, Kaplan SA. Serum 17-alpha-hydroxyprogesterone, progesterone, estradiol, and testosterone in the diagnosis and management of congenital adrenal hyperplasia. J Pediatr. 1974;85(6):782–787. doi:10.1016/s0022-3476(74)80340-9

14. Yushkevich PA, Piven J, Hazlett HC, et al. User-guided 3D active contour segmentation of anatomical structures: Significantly improved efficiency and reliability. NeuroImage. 2006;31(3):1116–1128. doi:10.1016/j.neuroimage.2006.01.015

15. Thomas R, Wang W, Su DM. Contributions of Age-Related Thymic Involution to Immunosenescence and Inflammaging. Immunity & Ageing. 2020;17(1):2. doi:10.1186/s12979-020-0173-8

16. Kooshesh KA, Foy BH, Sykes DB, Gustafsson K, Scadden DT. Health Consequences of Thymus Removal in Adults. N Engl J Med. 2023;389(5):406–417. doi:10.1056/NEJMoa2302892

17. Raptopoulou M, Goulis G. Physiological variations of T cells during the menstrual cycle. Clin Exp Immunol. 1977;28(3):458–460.

18. Lee S, Kim J, Jang B, et al. Fluctuation of peripheral blood T, B, and NK cells during a menstrual cycle of normal healthy women. J Immunol. 2010;185(1):756–762. doi:10.4049/jimmunol.0904192

19. Gaskins AJ, Wilchesky M, Mumford SL, et al. Endogenous reproductive hormones and C-reactive protein across the menstrual cycle: the BioCycle Study. Am J Epidemiol. 2012;175(5):423–431. doi:10.1093/aje/kwr343

20. Blum CA, Müller B, Huber P, et al. Low-grade inflammation and estimates of insulin resistance during the menstrual cycle in lean and overweight women. J Clin Endocrinol Metab. 2005;90(6):3230–3235. doi:10.1210/jc.2005-0231

21. Tracey KJ. The inflammatory reflex. Nature. 2002;420(6917):853–859. doi:10.1038/nature01321

22. Tracey KJ. Physiology and immunology of the cholinergic antiinflammatory pathway. J Clin Invest. 2007;117(2):289–296. doi:10.1172/JCI30555

23. Bassi GS, Kanashiro A, Coimbra NC, Terrando N, Maixner W, Ulloa L. Anatomical and clinical implications of vagal modulation of the spleen. Neurosci Biobehav Rev. 2020;112:363–373. doi:10.1016/j.neubiorev.2020.02.011

24. Réus GZ, Manosso LM, Quevedo J, Carvalho AF. Major depressive disorder as a neuro-immune disorder: Origin, mechanisms, and therapeutic opportunities. Neurosci Biobehav Rev. 2023;155:105425. doi:10.1016/j.neubiorev.2023.105425

25. Jarkas DA, Villeneuve AH, Daneshmend AZB, Villeneuve PJ, McQuaid RJ. Sex differences in the inflammation-depression link: A systematic review and meta-analysis. Brain Behav Immun. 2024;121:257–268. doi:10.1016/j.bbi.2024.07.037

26. Maes M, Berk M, Goehler L, et al. Depression and sickness behavior are Janus-faced responses to shared inflammatory pathways. BMC Med. 2012;10:66. doi:10.1186/1741-7015-10-66

27. Frank P, Jokela M, Batty GD, Cadar D, Steptoe A, Kivimäki M. Association Between Systemic Inflammation and Individual Symptoms of Depression: A Pooled Analysis of 15 Population-Based Cohort Studies. Am J Psychiatry. 2021;178(12):1107–1118. doi:10.1176/appi.ajp.2021.20121776

28. Ross JM, Barone JC, Tauseef H, et al. Predicting Acute Changes in Suicidal Ideation and Planning: A Longitudinal Study of Symptom Mediators and the Role of the Menstrual Cycle in Female Psychiatric Outpatients With Suicidality. Am J Psychiatry. 2024;181(1):57–67. doi:10.1176/appi.ajp.20230303

29. Okamura YT, Endo K, Toriihara A, et al. Automated quantitative evaluation of thymic involution and hyperplasia on plain chest CT. medRxiv. Preprint posted online November 14, 2023:2023.11.13.23298440. doi:10.1101/2023.11.13.23298440

